# HTSQualC is a Flexible and One-Step Quality Control Software for High-throughput Sequencing Data Analysis

**DOI:** 10.1101/2020.07.23.214536

**Authors:** Renesh Bedre, Carlos Avila, Kranthi Mandadi

**Affiliations:** Texas A&M AgriLife Research and Extension Center, Texas A&M University, Weslaco, TX, USA; Department of Horticultural Science, Texas A&M University, College Station, TX, USA; Department of Plant Pathology and Microbiology, Texas A&M University, College Station, TX, USA

**Keywords:** High-throughput sequencing (HTS), Quality control, Parallel computing, Software tool, CyVerse, Nextflow

## Abstract

Use of high-throughput sequencing (HTS) has become indispensable in life science research. Raw HTS data contains several sequencing artifacts, and as a first step it is imperative to remove the artifacts for reliable downstream bioinformatics analysis. Although there are multiple stand-alone tools available that can perform the various quality control steps separately, availability of an integrated tool that can allow one-step, automated quality control analysis of HTS datasets will significantly enhance handling large number of samples parallelly. Here, we developed HTSQualC, a stand-alone, flexible, and easy-to-use software for one-step quality control analysis of raw HTS data. HTSQualC can evaluate HTS data quality and perform filtering and trimming analysis in a single run. We evaluated the performance of HTSQualC for conducting batch analysis of HTS datasets with 322 samples with an average ∼1M (paired end) sequence reads per sample. HTSQualC accomplished the QC analysis in ∼3 hours in distributed mode and ∼31 hours in shared mode, thus underscoring its utility and robust performance. In addition to command-line execution, we integrated HTSQualC into the free, open-source, CyVerse cyberinfrastructure resource as a GUI interface, for wider access to experimental biologists who have limited computational resources and/or programming abilities.

## INTRODUCTION

Advancements in high throughput sequencing (HTS) technologies transformed biological research. HTS largely replaced conventional low-throughput Sanger-based sequencing technologies for genome-scale studies. Multiple genome sequencing approaches (DNA-seq, RAD-seq, GBS, AgSeq) are being used to study genetic variations, discovery of novel genes, high-throughput genotyping, biomarker discovery, and precision medicine ^1-5^. Similarly, transcriptome-level sequencing approaches (RNA-seq) allows determining the steady-state expression of genes, identification of novel transcripts and isoforms, alternatively splicing patterns, polymorphisms, gene co-expression networks, allele-specific expressions, and long non-coding RNAs (lincRNA) ^6-9^. Illumina’s benchtop and production-scale sequencers, which are by far the most widely used HTS platforms, can generate up to 1 and 20 billion sequence reads per run, respectively. This sequencing output is expected to rapidly increase due to further advancements in the Illumina technologies. As such advanced bioinformatics software tools are necessary to accelerate the analysis and efficiently manage the large volumes of data generated by the Illumina sequencing platforms.

In all HTS experiments, a critical first step is raw data quality assessment, filtering and trimming. HTS raw data often contains sequences of poor quality along with adapter or primer contaminations, and uncalled bases (N), which if not removed can significantly hamper the downstream bioinformatics analysis leading erroneous conclusions ^10-12^. Several standalone software tools are available for quality control analysis, trimming, and filtering of HTS data such as FastQC (https://www.bioinformatics.babraham.ac.uk/projects/fastqc/), NGS QC ^13^, FASTX-Toolkit (http://hannonlab.cshl.edu/fastx_toolkit/), NGS QCbox ^14^, Trimmomatic ^15^, fastp ^16^, and QC-Chain ^11^. However, most of them have limitations. For instance, FastQC performs only quality check of the data, and does not filter or trim of raw sequences (https://www.bioinformatics.babraham.ac.uk/projects/fastqc/). Conversely, FASTX-Toolkit although is equipped to perform quality filtering does not support parallel computing to handle large-scale batch analysis. Other software tools such as NGS QC, QC-Chain, Trimmomatic, and NGS QCbox have limited features for quality filtering, handles few samples at a time, dependent on other open-source software tools, and have a need to be run separately for different quality control features. Even though fastp has advantages over other tools, it does not support batch analysis ^16^. With HTS becoming more accessible and affordable, it is crucial to develop a flexible and integrative tool that can not only perform thorough quality control analysis, but can handle several hundred samples parallelly, with the least number of data handling steps.

Here, we present HTSQualC, which is an open-source and easy-to-use quality control analysis software tool for cleaning raw HTS datasets generated from Illumina sequencing platforms. HTSQualC is a flexible, one-step quality control software tool and can handle large number of samples. HTSQualC integrates filtering and trimming modules for single and paired end HTS data and supports parallel computing for batch quality control analysis. In addition to the quality filtering and trimming analysis, HTSQualC generates statistical summaries and visualization to assess the quality of the HTS datasets. HTSQualC can be used as command-line interface (CLI) as well as GUI. The GUI is available through CyVerse ^17,18^ Discovery Environment (https://cyverse.org/).

## MATERIALS AND METHODS

### Implementation

HTSQualC is a standalone open-source command-line software developed using Python 3 for quality control analysis of HTS data generated from Illumina sequencing platforms. The current HTSQualC version (v1.1) was developed specifically for Illumina generated FASTQ datasets, however, it could be utilized with other sequencing platforms given the input is FASTQ and supports same quality formats as Illumina. We focused on Illumina mainly because it is the most-widely utilized platform. We will continue development and subsequent versions of HTSQualC will be released to expand its use to other platforms such as PacBio and/or Nanopore Sequencing. At its core, HTSQualC consists of two main modules that are intended to filter and trim single and paired-end sequence datasets. Both modules were implemented using parallel computation to increase the performance of quality control analysis by allocating the input workload to the multiple CPUs. By default, there are only two CPUs, however this number can be changed as per user preferences. To some extent, HTSQualC works similar to MapReduce where it splits the large sequence file into smaller chunks, distributes the input data to multiple CPUs, and combines the input from each process to produce a final output.

HTSQualC checks quality issues in the raw HTS datasets and performs the quality filtering and/or trimming in a single run for removing low-quality bases, adapter contamination, and uncalled bases (N) as per the settings of the user. The entire quality control can be completed in a single input command. By default, HTSQualC filters out the sequence reads with Phred quality score < 20. We did not add the feature of duplicate reads removal in HTSQualC as this feature is directly related to the gene quantification and estimation of expression ^7^. A flowchart HTSQualC analysis is shown in Fig. 1.

**Fig. 1:**
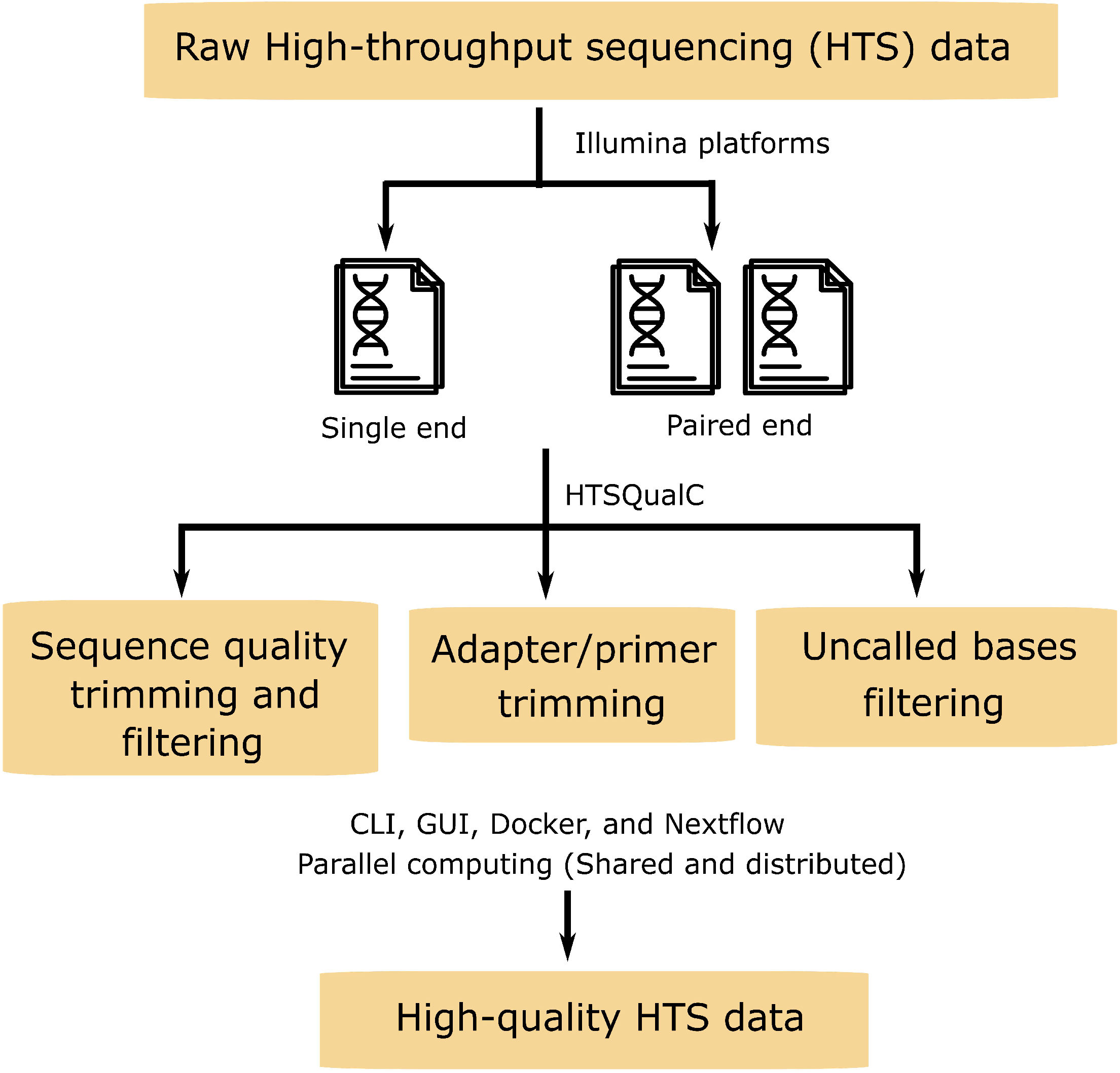
Flowchart of HTSQualC analysis. HTSQualC includes two main modules for quality control analysis of single and paired-end HTS datasets generated from Illumina sequencing platforms. HTSQualC filters and trims the raw HTS datasets to remove low-quality bases, adapter or primer contamination, and uncalled bases to generate high-quality datasets for downstream bioinformatics analysis.

HTSQualC can handle batch analysis of multiple sequencing datasets at a time. It accepts FASTQ file format as input and produces FASTQ or FASTA file format as output. HTSQualC accepts the GZ compressed FASTQ files and also provides an option to output GZ compressed FASTQ file. HTSQualC also generates summary statistics and visualization outputs for the filtered cleaned HTS datasets. All the outputs by default are saved in the same directory containing the raw RNA-seq input datasets. HTSQualC was primarily designed to run on the Linux and Mac operating systems as command-line interface CLI (Fig. 2), however, it can also run on the Windows operating systems using virtual machine. In addition, we made HTSQualC publicly available as GUI (Fig. 2) on CyVerse ^17,18^, which would be useful for experimental biologists with minimal bioinformatics training. CyVerse is an open-access, scalable, comprehensive, data analysis and management infrastructure for large-scale data analysis developed for life science research. CyVerse can be accessed through world-wide-web, and users can register for free at https://user.cyverse.org/register. To use the HTSQualC as GUI on CyVerse, it is necessary to have CyVerse account. Additionally, we have also provided the docker image and Nextflow template for running the HTSQualC.

**Fig. 2:**
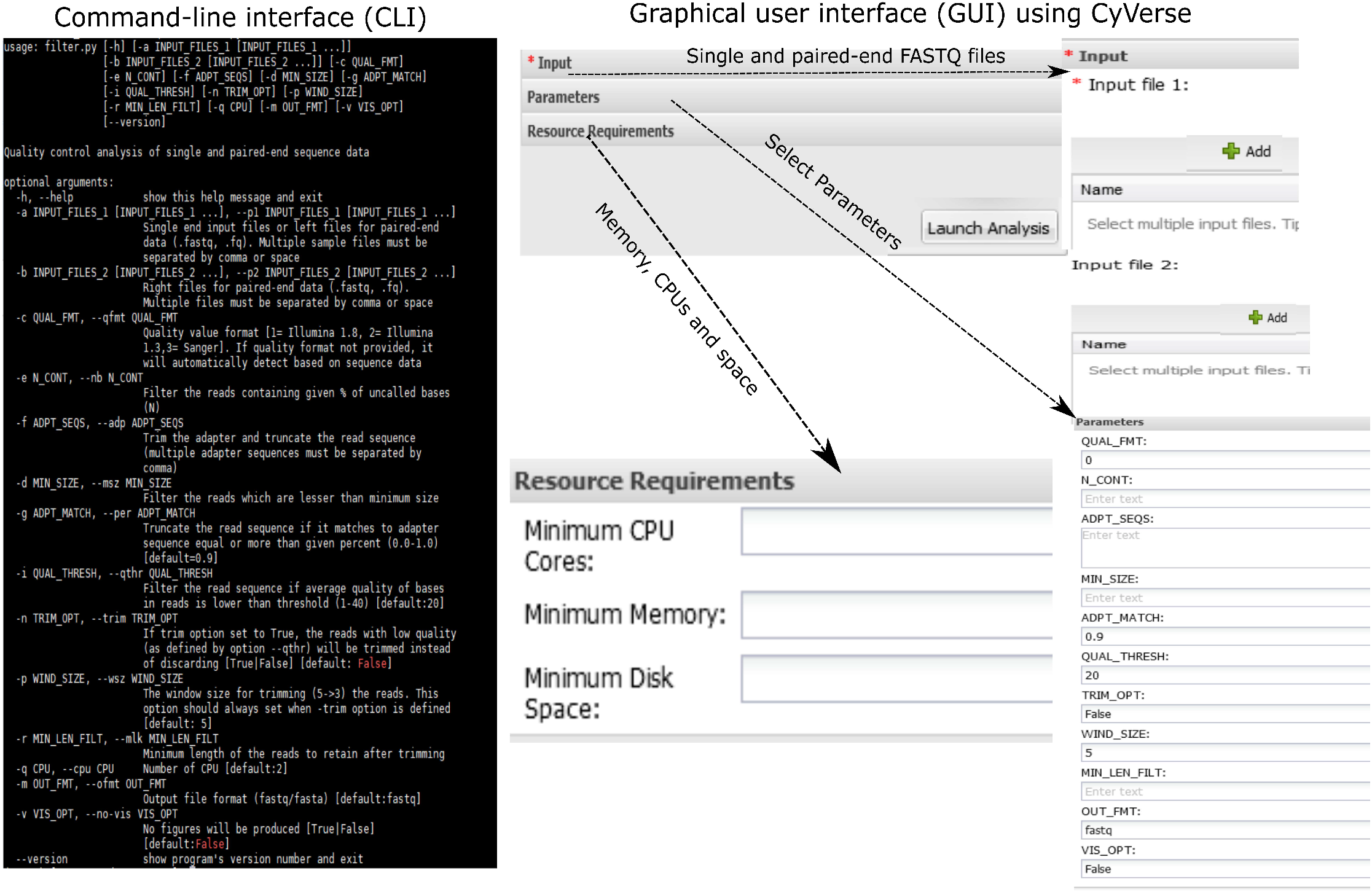
Command-line interface (CLI) and Graphic-user interface (GUI) of HTSQualC. HTSQualC can be launched as CLI or GUI modules. For CLI, the HTSQualC need to installed on a local system, whereas GUI is preinstalled and ready to use online at CyVerse (https://cyverse.org/).

## RESULTS AND DISCUSSION

### Case studies

To evaluate the performance of HTSQualC for quality control analysis, we analyzed several datasets using a 64 GB RAM and 20 CPUs computing node (Intel 2.5 GHz IvyBridge processor) of Texas A&M High Performance Research Computing Center (HPRC) (http://hprc.tamu.edu/). First, we analyzed a single-end raw RNA-seq dataset generated on Illumina HiSeq 2000 platform corresponding to cotton, a dicot plant (BioProject accession PRJNA275482 and SRA ID: SRR1805340, Table 1) ^19,20^. HTSQualC analysis was performed with the default parameters. HTSQualC automatically detected the Illumina sequence quality variant and filtered out the reads with Phred quality score < 20. In total, HTSQualC filtered out ∼5 M reads (Fig.3 and Supplementary File 1A).

**Table 1:** Summary of the quality control analysis of single and paired-end datasets with single and multiple samples performed using HTSQualC CLI.

| SRA Accession or Samples | Read Type (# Samples) | # Sequence reads (File size in GB<sup>a</sup>) | Read length (bp) | # CPUs (Parallel computing) | Run time (Min) |
| --- | --- | --- | --- | --- | --- |
| SRR2165176 | Paired (1) | 8,583,424×2 (5.6) | 100 | 18 (Shared) | 9 |
| SRR2165176 | Paired (3) | 8,583,424×2 (5.6) | 100 | 18 (Shared) | 28 |
| SRR2165177 |  | 9,282,222×2 (6) | 100 |  |  |
| SRR2165178 |  | 9,918,081×2 (6.4) | 100 |  |  |
| SRR1805340 | Single (1) | 82,059,811 (27) | 100 | 18 (Shared) | 40 |
| Tomato GBS<sup>b</sup> | Paired (322) | ~ 1,000,000×2 (206) | 150 | 18 (Shared) | 1855 |
|  |  |  |  | 18 (Distributed) | 157 |

<sup>a</sup>Combined file size of two files for paired-end sequence reads
<sup>b</sup>322 tomato genotypes were sequenced using low-coverage whole genome sequencing by Illumina HiSeq 4000 <sup>5</sup>.
The number of sequences reads are average of all the datasets.

**Fig. 3:**
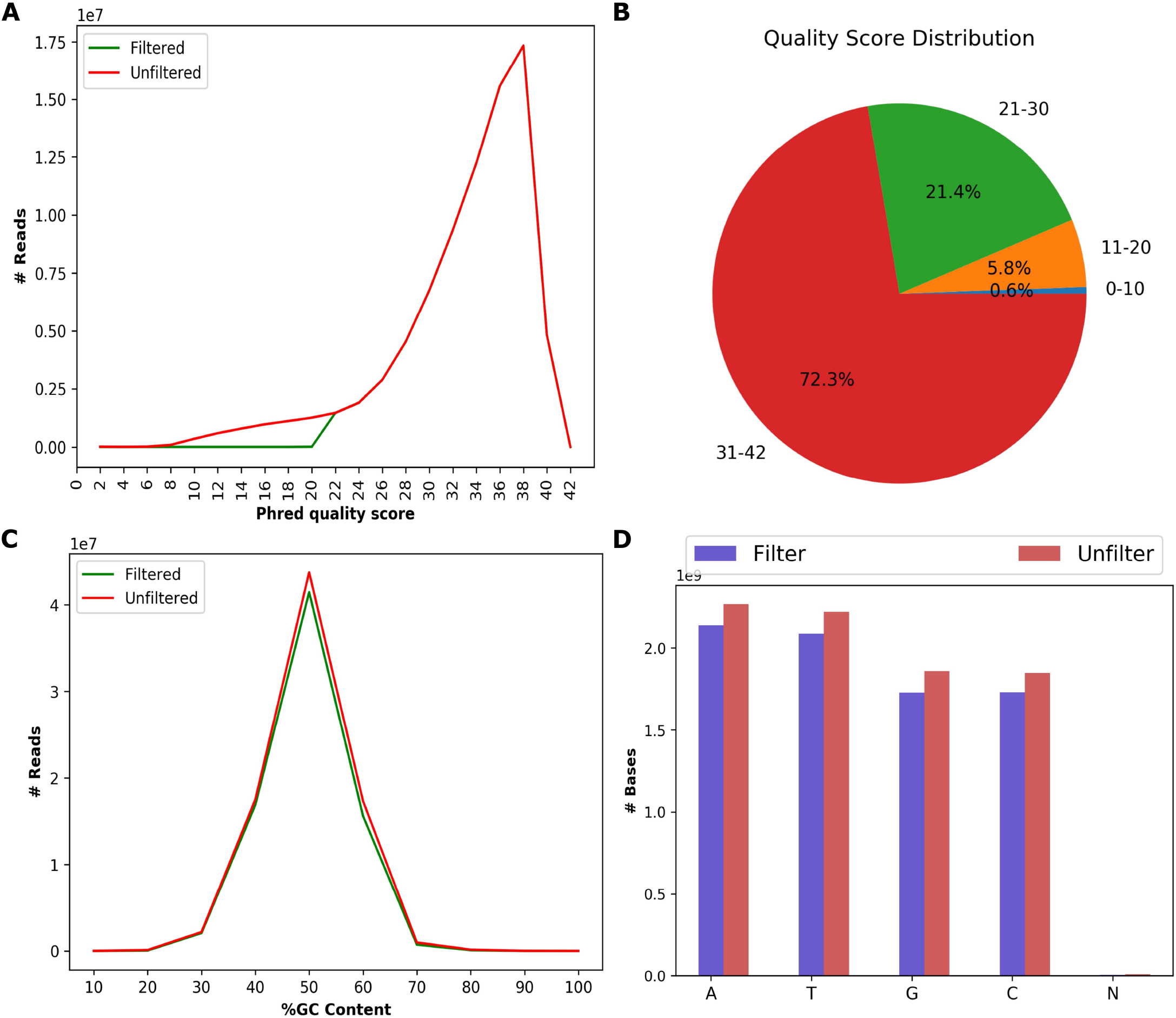
The sequence quality evalaution of HTS datatsets performed by HTSQualC. A) The sequence quality distribution among the raw (unfiltered) and cleaned (filtered) sequence data, B) The distribution of percenatges of sequence reads with quality score, C) The percentage GC content distribution among the raw and cleaned sequence data, D) The content of nucleotide bases in raw and cleaned sequence data.

Second, we evaluated multiple paired-end raw RNA-seq datasets generated on Illumina HiScanSQ platform corresponding to sugarcane, a monocot plant (BioProject accession PRJNA291816 and SRA IDs: SRR2165176, SRR2165177, SRR2165178) ^6,21^. Initially, we ran HTSQualC on one paired-end dataset (SRR2165176) using default parameters (Table 1). HTSQualC was able to analyze the Illumina sequence quality variants and filtered out the reads with Phred quality score < 20 (Supplementary File 1B). For instance, the HTSQualC filtered out the ∼250 K sequence reads which were below the quality threshold (Supplementary File 1B). Next, we ran the HTSQualC with customized parameters for filtering adapter sequences, quality thresholds, and uncalled bases. In total, HTSQualC filtered out ∼451 K reads and trimmed ∼20 K reads (Supplementary File 1C). All the HTSQualC commands used for performing the above analyses are provided in the README file.

In addition to quality control analysis, we also evaluated the processing time and batch handling of HTSQualC using default parameters. HTSQualC took ∼9 min to perform the quality filtering analysis of ∼9 M paired-end sequence reads, and ∼40 min for ∼82 M single-end sequence reads with 18 CPUs (Table 1).

Lastly, we analyzed 322 paired-end genotyping-by-sequencing (GBS) datasets corresponding to tomato ^5^. These datasets had ∼1M (×2) sequence reads per sample. For this experiment, we also compared parallel computing in shared vs. distributed mode with 18 CPUs per computing node using Nextflow. We used a single HPRC node for shared mode and several HPRC nodes for distributed mode. We were able to analyze the 322 datasets in ∼1855 min (∼31 hours) and ∼157 min (∼3 hours) in the shared and distributed computing modes, respectively (Table 1).

### Advantages and disadvantages of HTSQualC over existing quality control analysis software

We compared the advantages of HTSQualC with prevailing quality control analysis tools such as the FastQC (https://www.bioinformatics.babraham.ac.uk/projects/fastqc/), NGS QC ^13^, QC Chain ^11^, FASTX-Toolkit (http://hannonlab.cshl.edu/fastx_toolkit/), fastp ^16^ and NGS QCbox ^14^ on various important quality control features (Table 2). Many of the current tools are not as integrated as HTSQualC (Table 2).

**Table 2:** The key sequence quality features comparison of HTSQualC with other leading and equivalent quality control software tools.

| Features | HTSQualC | FastQC<sup>a</sup> | NGS QC | QC-Chain | FASTX-Toolkit | NGS QCbox | fastp |
| --- | --- | --- | --- | --- | --- | --- | --- |
| Low quality filtering | Yes | Yes | Yes | Yes | Yes | No | Yes |
| Uncalled bases (N) | Yes | No | Yes* | No | No | No | No |
| Adapter or primer trimming | Yes | Yes | Yes | Yes | Yes | No | Yes |
| Multiple adapter support | Yes | Yes | Yes | Yes | No | No | Yes |
| Minimum read length filtering | Yes | No | No | No | No | Yes | Yes |
| Low quality trimming | Yes | No | Yes* | Yes | No | Yes | Yes |
| Paired-end support | Yes | No | Yes | Yes | No | Yes | Yes |
| Paired-end sequence order | Yes | No | Yes | Yes | No | Yes | Yes |
| Visualization | Yes | Yes | Yes | Yes | Yes | Yes |  |
| Parallel computing | Yes | Yes | Yes | Yes | No | Yes | Yes |
| Automatic quality variant detection | Yes | No | Yes | No | No | No | No |
| FASTA Output | Yes | No | No | No | No | No | No |
| Programming | Python 3 | Java | PERL | C++ | C | C | C/C++ |
| GZIP FASTQ input | Yes | No | Yes | No | No | Yes | Yes |
| Multiple sample support (batch analysis) | Yes | Yes | Yes | No | No | Yes | No |
| GitHub | Yes | Yes | No | No | Yes | Yes | Yes |

<sup>a</sup>FastQC only provides quality metric and does not filter or trim HTS data
\*Separate tools for given features

For instance, programs such as FastQC only provides the quality control statistics of the raw HTS data and cannot be used for quality filtering. The FASTX-Toolkit does quality filtering, but the various trimming, filtering options are not integrated, and each analysis must be performed separately. The NGS QCbox comes close, however only allows sequence read trimming based on a quality threshold for specific window size and it is highly dependent on other open-source software tools for quality control analysis ^14^. NGS QC allows filtering and trimming of low-quality reads similar to FASTX-Toolkit, however these software tools are independent and must be run separately. The fastp provides single run quality control analysis of FASTQ files similar to HTSQualC but does not offer key features such as batch quality control analysis, automatic sequence quality format detection, etc. The fastp does offer other features which are not included in HTSQualC such as poly tail trimming, UMI preprocessing, and basic support for the long-read sequence data. Hence, in this context, the two tools can be complementary to each other for a range of applications.

The removal of sequence reads containing uncalled bases (N) is critical to improve the accuracy of the sequence reads ^22^. HTSQualC has this feature of filtering the sequence reads based on the content of the uncalled bases (N). Although NGS QC allows this analysis, it is not integrated, and must be run separately ^13^. In addition to overcoming several of these limitations, HTSQualC supports common features with NGS QC and NGS QCbox such as allowing inputs in the compressed GZIP file format for seamless input of large datasets (Table 2). Furthermore, similar to NGS QC, HTSQualC has an integrated function to automatically detect the FASTQ quality variants (Table 2). Only HTSQualC provides a parameter to output the cleaned FASTQ data in FASTA format, avoiding using additional file format conversion tools. Lastly, in addition to the command line execution, we integrated HTSQualC as a graphic user interface that is accessible freely for everyone to access through CyVerse.

We compared the running times of HTSQualC with FastQC, FASTX-Toolkit, and fastp with default settings for quality filtering Among these three tools, fastp took the least amount of time (2 minutes), followed by FastQC (3 minutes) and HTSQualC (24 minutes). The HTSQualC took longer as it is developed in Python 3 (interpreted language), which is slower than C/C++ and Java, while FASTX-Toolkit took the longest time (33 minutes) mainly because it does not support parallel computation. When we compared the memory consumption, the fastp (∼420 MB) and HTSQualC (∼818 MB) outperformed FastQC (1.55 GB). Because HTSQualC works in the same way as MapReduce, it will require more storage because of the numerous input and output processes. Furthermore, as HTSQualC is nearing completion, it automatically deletes the intermediate files, reducing further manual steps required to clear the intermediate files. We could not evaluate NGS QC toolkit, QC chain, and NGS QCbox as these tools were not available or currently not maintained anymore.

We also evaluated the output of HTSQualC and fastp with default settings for quality filtering. For quality filtering, HTSQualC and fastp have different algorithmic implementations. For instance, by default, fastp performs the quality filtering on Phred quality score and discards the sequencing reads where certain percentages of bases have Phred quality <15 (<97% base call accuracy), whereas HTSQualC discards the reads which have average Phred quality <20 (<99% base call accuracy). With default settings, fastp filtered out ∼115K reads, whereas HTSQualC filtered out 4256 reads on an Illumina dataset containing ∼25 M reads. It is advisable to keep the base call with high Phred quality (>20 or >30) to minimize calling false-positive variant calls ^23^. HTSQualC also offers more data outputs, such as the sequence quality format, minimum, maximum, and mean read lengths, and average Phred quality values of filtered and unfiltered reads, which are not included in the fastp output (Supplementary File 2). The output details are provided in Supplementary File 2.

## CONCLUSION

HTSQualC is an open-source, integrated, and easy-to-use software designed for one-step quality control visualization and quality filtering of raw HTS data generated using the Illumina sequencing platforms. The flexiblility to detect and remove the low-quality sequences, adapter or primer contamination, uncalled bases, in a single run, greatly enhances the automation of HTS data analysis projects. Because HTSQualC can be implemented by parallel computing, it enables batch handling of large number (>300) of datasets. In addition to the command line interface, HTSQualC is available as graphic user interface that is accessible freely through CyVerse. The later should significantly facilitate its use among biologists without much prior bioinformatics or command line computing experience.

## Acknowledgements

We thank Upendra Devisetty, Reetu Tuteja, and Sarah Roberts for their assistance with installing HTSQualC at CyVerse DE, which was made possible through CyVerse’s External Collaborative Partnership program. We acknowledge support of Texas A&M High Performance Research Computing Center (http://hprc.tamu.edu/) resources and sequencing support of the Texas A&M AgriLife Genomics and Bioinformatics Service (https://txgen.tamu.edu/). We also acknowledge support of Louisiana State University Agricultural Center and Louisiana State University High Performance Computing resources (http://www.hpc.lsu.edu/) for supporting early stages of the HTSQualC development as part of Ph.D. research of RB. This work was supported in part by funds from Texas A&M AgriLife Research Insect-vectored Disease Seed Grant (114190-96210) to KM, Foundation for Food and Agricultural Research New Innovator Award (2018-534299) and USDA-NIFA (2018-70016-28198, HATCH 1023984) awards to KM.

## Author contributions

RB developed the software, platform and conducted the analysis. CA, and KM supervised the study, data analysis and interpretation. All authors have read, reviewed, and approved the manuscript.

## Availability

HTSQualC software, Docker image and Nextflow template are available for download at https://github.com/reneshbedre/HTSQualC and graphical user interface (GUI) is available at CyVerse Discovery Environment (DE) (https://cyverse.org/). Documentation is available at https://reneshbedre.github.io/blog/htseqqc.html and https://cyverse-htseqqc-cyverse-htseqqc-cyverse-tutorial.readthedocs-hosted.com/en/latest/. The HTSQualC is also available on Anaconda cloud (https://anaconda.org/bioconda/htsqualc) and can be installed using biconda channel.

